# Expression and fusogenic activity of SARS CoV-2 Spike protein displayed in the HSV-1 Virion

**DOI:** 10.1101/2023.11.28.568860

**Authors:** Prashant J. Desai

## Abstract

Severe acute respiratory syndrome coronavirus (SARS-CoV) is a zoonotic pathogen that can cause severe respiratory disease in humans. The new SARS-CoV-2 is the cause of the current global pandemic termed coronavirus disease 2019 (COVID-19) that has resulted in many millions of deaths world-wide. The virus is a member of the Betacoronavirus family, its genome is a positive strand RNA molecule that encodes for many genes which are required for virus genome replication as well as for structural proteins that are required for virion assembly and maturation. A key determinant of this virus is the Spike (S) protein embedded in the virion membrane and mediates attachment of the virus to the receptor (ACE2). This protein also is required for cell-cell fusion (syncytia) that is an important pathogenic determinant. We have developed a pseudotyped herpes simplex virus type 1 (HSV-1) recombinant virus expressing S protein in the virion envelop. This virus has also been modified to express a Venus fluorescent protein fusion to VP16, a virion protein of HSV-1. The virus expressing Spike can enter cells and generates large multi-nucleated syncytia which are evident by the Venus fluorescence. The HSV-1 recombinant virus is genetically stable and virus amplification can be easily done by infecting cells. This recombinant virus provides a reproducible platform for Spike function analysis and thus <u>adds</u> to the repertoire of pseudotyped viruses expressing Spike.

**Impact Statement:** The isolation of a pseudotyped herpes simplex virus type 1 (HSV-1) virus using the Spike protein is new and innovative. This virus can be used to study entry and fusion events mediated by the S protein as well as test antibodies for their ability to neutralize this particle. In addition, these virions can be used for screening antibody specificity using the S protein displayed in its natural membrane bound conformation.

---

Coronaviruses are enveloped, non-segmented, positive-sense RNA viruses that carry a ∼30,000 nucleotide genome [1–4]. The spherical structural particle of the virus is about 80-125 nm in diameter and the virus envelop is embedded with three membrane proteins: the spike (S) protein that gives the virus the “corona” structure [5], and the envelope (E) and membrane (M) proteins [1, 2]. The virus membrane envelops the RNA genome which is encapsidated by the nucleocapsid (N) protein. The spike protein engages the angiotensin converting enzyme-2 (ACE2) receptor [6, 7] and acts to fuse viral and cellular membranes during entry [8, 9]. The E protein plays roles in virus assembly and budding but also has other roles. The M protein is the most abundant envelop protein and coordinates virus assembly and budding through protein-protein interactions with the other virion components [10, 11]. The N protein with the viral genome form the ribonucleoprotein core and has been shown to be involved in viral RNA synthesis, transcriptional regulation of genomic RNA, translation of viral proteins, and budding [3]. The Spike protein is the structural protein responsible for the crown-like shape of the CoV viral particles because it forms a trimeric complex. The 1255 aa long protein (∼185 kD glycosylated polypeptide) is a class-I viral fusion protein [12] and contributes to the cell receptor binding, tissue tropism and pathogenesis. The Spike protein is cleaved by host cell proteases at the S1/S2 cleavage site [5, 13–15]. Following cleavage, also known as priming, the protein is divided into an N-terminal S1-ectodomain that recognizes the cognate cell surface receptor and a C-terminal S2-membrane-anchored protein involved in viral entry by membrane fusion. The S1-protein contains a conserved Receptor Binding Domain (RBD), which recognizes the angiotensin-converting enzyme 2 (ACE2) receptor [16]. Cell-cell fusion resulting in syncytia formation (multi-nucleated cell) is a characteristic property of the Spike protein [17–19]. Syncytia formation also likely contributes to the pathology of the disease as observed by the presence of multinucleate pneumocytes in patients with advanced disease [20–25]. The biological processes that define Spike protein in the virion binding to the cell (entry) and Spike protein in infected cell membrane fusing with uninfected cells (syncytia formation) are similar in that they both require binding to ACE2 receptor and proteolytic activation to expose the fusion peptide [18, 26–29]. However, differences between these two processes are evident and the means to inhibit them may similarly differ [30, 31]. Cell-cell transmission of the virus is also a means by which the virus can evade some neutralizing antibodies [31, 32].

Our goal was to leverage our expertise in membrane protein display and self-assembly of virion structures to develop and create tractable models to study this highly pathogenic virus. Our first goal was to use herpes simplex virus type 1 (HSV-1) to express and display the membrane proteins of SARS-CoV-2. Use of HSV-1 to display Spike protein could be useful for investigation for serology, monoclonal antibody screening/specificity testing and as pseudotyped HSV-1 virions that can be used safely to examine entry inhibition and virus neutralization. The pseudotyped virus is also genetically stable, does not have to be re-made each time by transfection methods and can be used safely in a BSL2 level facility.

Previously, we have expressed and displayed membrane proteins to provide novel platforms and tools for investigation of their functional activities using the Virion Display (VirD) method [33–35]. HSV-1 produces large spherical virions displaying hundreds of copies of envelop proteins. Our aim was to engineer this virus to express human membrane proteins during the virus productive cycle and incorporate the human proteins into the virion during the assembly process. This was achieved by cloning the membrane protein gene in place of the glycoprotein B gene of HSV-1 (UL27) such that the expression of the human membrane protein is driven by the gB (UL27) promoter. Because the gene is now expressed as a “viral” gene it was subsequently incorporated into the virion envelop during virus assembly. The expression of the human membrane proteins in infected cells, at the cell surface and in purified virions, was in the correct transmembrane orientation, and the proteins are biochemically functional [33, 34]. Subsequently, we engineered the HSV-1 genome to be Gateway compatible by inserting the Gateway selection cassette in the UL27 gene locus. This locus encodes glycoprotein B of HSV-1 which is the major mediator of cell fusion. Membrane protein ORFs can be cloned into this site using standard Gateway cloning methods [34].

We obtained a codon optimized Spike protein ORF from BEI resources (WuHan strain). Plasmid pCAGGS was used as a template to PCR amplify the ORF using Q5 (NEB) polymerase. The primers contained the attB recombination sequences compatible with Gateway cloning. A BP reaction was performed using BP clonase (Invitrogen) and pDONR221. Transformants were screened for the recombined ORF and four clones were sequenced. These four validated clones were used to transfer the Spike ORF into the HSV-1 strain KOS bacmid using LR clonase (Invitrogen) to derive four recombinant viruses S1-S4. The HSV-1 bacmid carries a Gateway cassette such that the expression of the cloned membrane protein is driven by the gB (UL27) promoter and the C-terminus is tagged with a V5 epitope sequence so we can monitor the expression of the protein. Gateway cloning methodology has been described in more detail in Syu *et al*. [34]. Reconstitution of infectivity of the HSV-1 bacmid was performed as previously described using the gB complementing cell line, D87 [34]. D87 is a Vero cell line that expresses gB upon superinfection. All cell lines and virus stocks were prepared as described by Desai *et al.* [36].

To examine expression of the Spike protein, Vero cells (5 x 10^5^) were infected at a multiplicity of infection (MOI) of 10 plaque forming units (PFU)/cell. The infected cells were harvested at 24 hour post-infection and protein lysates prepared in Laemmli buffer. Proteins were analyzed on 4-12% NuPage gels (Invitrogen) and transferred to nitrocellulose membranes using the iBlot instrument (Invitrogen) as previously described [37]. The blots were reacted with mouse anti-V5 antibody (Invitrogen). Abundant quantities of the Spike protein were observed in the lysates of all four isolates (Fig. 1). The Spike protein is 1255 amino acids long and is predicted to have a molecular weight of approximately 185 kD (glycosylated). We also observed a proteolytic cleaved product that is V5 reactive.

**Fig. 1.**
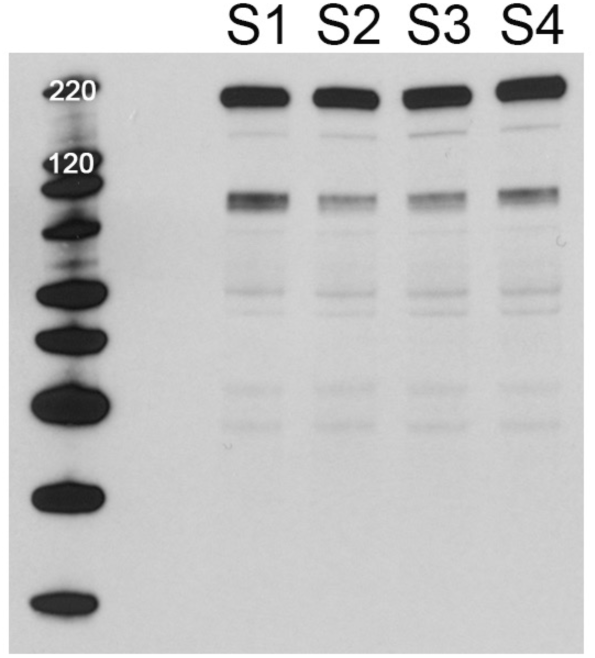
Expression of SARS-CoV-2 Spike protein in the HSV-1 KOS strain. The Spike protein encoding gene was cloned into the UL27 locus of HSV-1. The ORF is fused at the C-terminus with a V5 epitope tag. Four isolates were purified and amplified for investigation. Cells were infected with all four viruses and harvested 24h after infection. The protein lysates were examined for Spike protein expression using immunoblot methods and V5 antibody. Protein with the correct mobility was detected in all four lysates. In addition, a cleavage product was also present in each lysate and of approximately 90 kD in size. Protein standards are in the left lane.

To examine whether the Spike protein was incorporated into HSV-1 virions, Vero cells (1 x 10^7^) were infected with the recombinant viruses at an MOI of 10 PFU/cell. The supernatants of the infected cells were collected at 24 hours post-infection and layered onto a 20% sucrose cushion. This was centrifuged at 39 K for 20 minutes in a Beckman SW41 rotor. The virion pellet was resuspended in Laemmli buffer and analyzed by immunoblot as described above. Immunoblot data show the Spike protein is incorporated into HSV-1 extracellular virions (Fig. 2). Both in the infected cells and more so in the virion we observed a cleavage product that is V5 tagged (Fig 2). We assume this is the S2 polypeptide and it has a mobility around 90 kD. SARS-CoV-2 Spike protein is proteolytically cleaved first by furin at the S1/S2 site and subsequently at the S’ position by serine protease 2, TMPRSS2 [38, 39]. In Vero cells, furin is likely the major processing enzyme although endogenous levels of TMPRSS2 may also cleave the Spike protein [14]. Our analysis cannot resolve that with the HSV-1 Spike protein expression seen in Fig. 2, however, the cleavage that we observed was functional for syncytia formation. Mutations in the polybasic cleavage site have been observed upon passage of the virus in culture (Vero cells) and there is debate about role of the furin cleavage site for infectivity [28, 40–42].

**Fig. 2.**
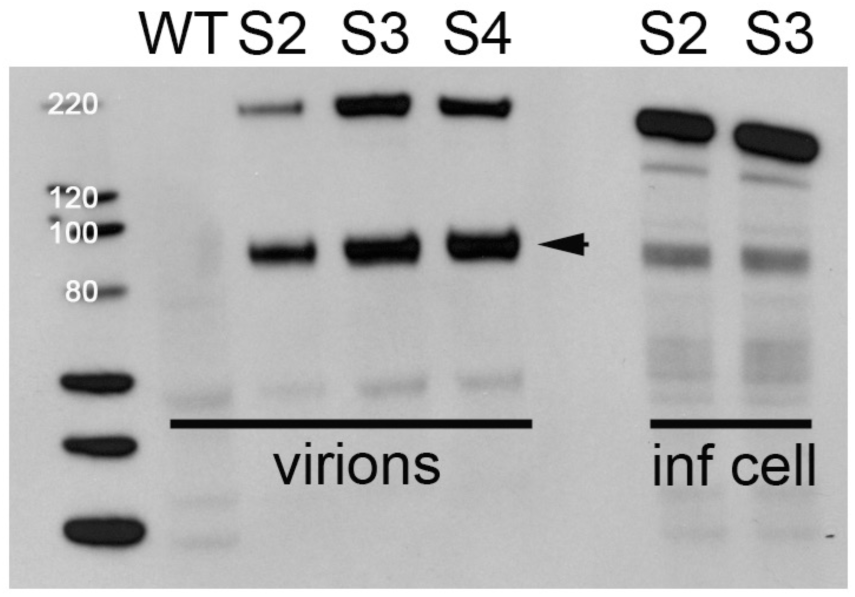
Incorporation of the SARS-CoV-2 Spike protein in HSV-1 virions. Virions in the extracellular media were collected and purified through a sucrose cushion. The virions were then analyzed by immunoblot methods using V5 antibody. Infected cell lysates were included as positive controls. S2, S3 and S4 virions all contained Spike protein Wild-type (WT) virions do not. A cleavage product (arrow) with a mobility of approximately 90 kD was also present in the virions. Protein standards are in the left lane.

We also recombined into the S3 virus, a VP16-Venus marker which allows one to follow HSV-1 virus entry and replication [43]. VP16 is a major virion (tegument) protein of HSV-1. VP16 localizes to nuclear puncta and then subsequently also at cytoplasmic sites during the infectious cycle. Recombination was done by co-infecting D87 cells with the S3 virus and a virus that expresses VP16-Venus as well as mutations in UL16 and UL21 that are lethal [43]. The virus from the co-infected cells was plaqued on D87 cells. Plaques that formed on D87 cells that were also fluorescent were purified further. Subsequently an S3 virus expressing VP16-Venus was amplified and used in the experiments below.

In the recombinant HSV-1 virus, the cloned gene is in the UL27 locus which encodes glycoprotein B (gB) [35]. This HSV-1 recombinant cannot replicate because of the loss of the essential UL27 gene [44, 45]. Glycoprotein B of HSV-1 is the major fusogen for HSV-1 and facilitates fusion of virion and cell membranes following receptor binding [46, 47]. Thus, we also investigated whether the SARS Spike protein can provide this function to HSV-1 virions that cannot fuse and enter, essentially pseudotyping HSV-1. We took virions that lack gB but now have Spike protein in their membranes and infected Vero cells. We observed VP16-Venus fluorescence in single cells indicative of virus entry. These initial fluorescent puncta progressed to large syncytia as time passed (Fig. 3A). Syncytia formation in cells infected by these viruses indicated cell-cell fusion mediated by S protein, was also functional in our assay.

**Fig. 3.**
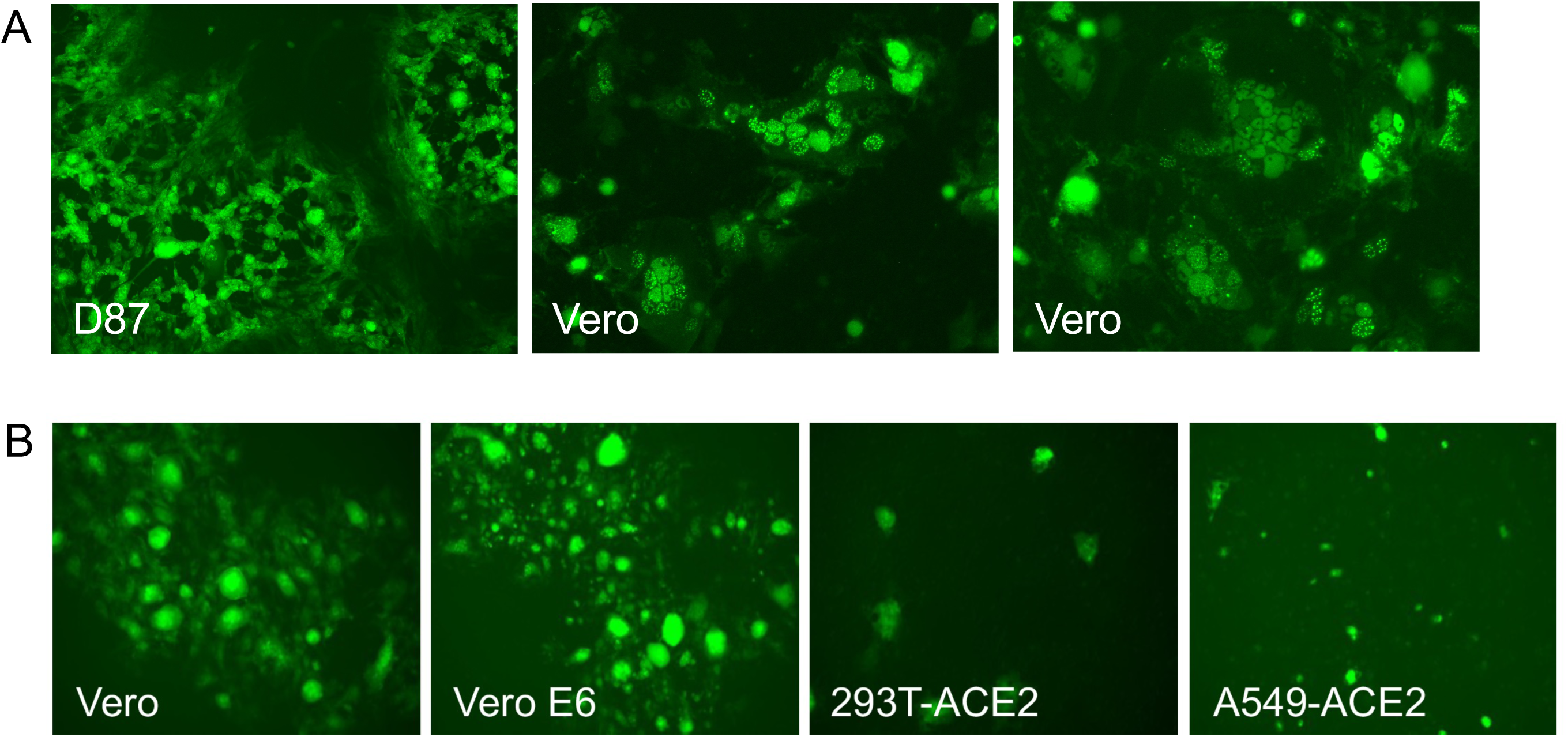
Panel A. HSV-1 fluorescent tagged virus pseudotyped with Spike protein. A VP16-Venus fluorescent fusion protein tag was incorporated into the genome of the S3 HSV-1 virus. Virus was plaqued on the complementing D87 cell line and visualized by a fluorescence microscope. This virus following infection of Vero cells formed extensive syncytia indicative of cell-cell fusion. Panel B. Syncytia formation in different cell lines. Cells were infected with HSV-1 pseudotyped with Spike protein. Imaging was done by fluorescence microscopy using a Zoe microscope (BioRad) using the X10 objective lens.

We compared different cell lines to examine syncytium formation, Fig. 3B. Vero cells are typically used for propagation and plaquing of the HSV-1 Spike virus. We also used two cell lines transduced with the ACE2 expression vector, HEK-293T and a human lung cell line A549 [48]. Vero and Vero E6 cells were the best cell lines for visualization of cell-cell fusion. What was astonishing is the size of the syncytia formed in these cells. Therefore, this was a fortuitous tag that visually illuminates the syncytium.

It has been reported that the HIV protease inhibitor, nelfinavir can inhibit cell fusion mediated by the Spike protein [49]. We also tested this using the recombinant virus S3-Venus. At concentrations of 15 and 20 uM, nelfinavir inhibited cell fusion mediated by the Spike protein (data not shown).

The major goals of this investigation was to establish a pseudovirus system for SARS CoV-2 that offers a safe tool to study virus entry and membrane fusion. The isolation of a pseudotyped HSV-1 virus using the Spike protein is innovative. Futhermore, this reagent provides a powerful platform to elucidate antibody responses, functional activities, identify neutralizing monoclonal antibodies and therapeutic approaches. We have already demonstrated we can generate S protein displaying virions and they have biological activity in that they support virus entry and extensive cell-cell fusion. In addition, these virions can be used for screening antibody specificity using the S protein displayed in its natural membrane bound conformation. While the Knipe Lab made an HSV-1 recombinant expressing S protein, theirs was a replication defective mutant and thus does not produce virions [50]. Their goals were to investigate host responses to S protein expressed in different human cell lines.The pseudotyped virus is also genetically stable, does not have to be re-made each time by transfection methods, unlike the lentivirus based methods, and can be used safely in a BSL2 facility.

## Acknowledgements

BEI resources provided plasmids and cell lines. We thank Andrew Pekosz (Johns Hopkins School of Public Health) for Vero E6 cells.

